# B cell directed CAR-T cell therapy results in activation of CD8+ cytotoxic CAR-negative bystander T cells in both non-human primates and patients

**DOI:** 10.1101/2023.09.28.559936

**Authors:** James Kaminski, Ryan A. Fleming, Francesca Alvarez-Calderon, Connor McGuckin, Emily E. Ho, Fay Eng, Xianliang Rui, Paula Keskula, Lorenzo Cagnin, Joanne Charles, Jillian Zavistaski, Steven P. Margossian, Malika A. Kapadia, James B. Rottman, Jennifer Lane, Susanne H.C. Baumeister, Victor Tkachev, Alex K. Shalek, Leslie S. Kean, Ulrike Gerdemann

**Affiliations:** Division of Pediatric Hematology-Oncology, Boston Children’s Hospital, Boston, MA; Broad Institute of MIT and Harvard, Cambridge, MA, United States; Department of Pediatric Oncology, Dana-Farber Cancer Institute, Boston, MA; Harvard Medical School, Boston, MA; 2seventy bio, Cambridge, MA; Center for Transplantation Sciences, Massachusetts General Hospital, Boston, MA 02129; Department of Chemistry, Institute for Medical Engineering and Science (IMES), and Koch Institute for Integrative Cancer Research, MIT, Cambridge, MA, United States; Ragon Institute of Massachusetts General Hospital (MGH), MIT and Harvard, Cambridge, MA, United States

## Abstract

There is growing appreciation for the emergence of CAR^neg^ bystander T cells after CAR-T cell infusion. However, their phenotypic and transcriptomic hallmarks and mechanisms of activation remain uncertain. We performed single-cell RNA-Seq (scRNA-Seq) on non-human primate (NHP) and patient-derived T cells to interrogate CAR^neg^ T cells following B cell targeted CAR-T cell therapy. In a NHP model, we observed a distinct population of activated CD8+ CAR^neg^ T cells emerging during CAR-T cell expansion. These bystander CD8+ CAR^neg^ T cells exhibited a unique transcriptional signature with upregulation of NK-cell markers (*KIR3DL2, CD160, KLRD1*), chemokines and chemokine receptors (*CCL5, XCL1, CCR9*), and downregulation of naive T cell-associated genes (*SELL, CD28*). A transcriptionally similar population was identified in patients following Tisangelecleucel infusion. Mechanistic studies revealed that IL-2 and IL-15 exposure induced bystander-like CD8+ T cells. These T cells efficiently killed leukemic cells through a TCR-independent mechanism. Together, these data identify bystander CD8+ T cells as a novel mechanism by which CAR-T cell infusion can induce further anti-leukemic activity, measurable in both NHP and in patients.

**Statement of Significance:** We have deeply interrogated CAR^neg^ bystander CD8+ T cells during CAR-T cell expansion in non­human primates and patients receiving Tisangelecleucel to identify the unique transcriptomic signature defining these cells, and to determine that IL-2-and IL-15-induced cytotoxic bystander T cells are capable of killing in a TCR-independent manner. These data highlight the potential of bystander T cells for leukemia control and provide a critical foundation for their future analysis.

## Introduction

Chimeric antigen receptor T cells (CAR-T cells) are a breakthrough therapy, capable of inducing remission in patients with relapsed or refractory B cell-derived malignancies, including both B-lineage leukemias and lymphomas.^1–4^ However, despite high rates of initial response, some patients are refractory to CAR-T cells, and up to 50% of responding patients will eventually experience relapse.^3,4^ As experience with CAR-T cells has grown, the field has gained an increased understanding of the factors that determine the success or failure of these cells,^1,5^ with several studies using single-cell RNA-Seq (scRNA-Seq) to identify key transcriptional drivers of CAR-T cell function.^6–12^ Unfortunately, despite the growing biological insights into CAR-T cell profiles and their association with disease control, it is still not possible to accurately predict which patients will respond and which patients will be resistant to CAR-T cell treatment. An additional aspect of the therapeutic effect of CAR-T cells, which is now gaining increasing attention, is their impact on surrounding immune cells,^13^ and the potential of these ‘bystander’ cells to also elicit an anti-tumor response.^13^

The phenomenon of bystander activation was initially described in viral infections, where T cells lacking TCRs cognate to viral antigens were discovered to contribute to viral clearance.^14^ Recent studies have found evidence of bystander activated T cells within the tumor (including B cell lymphoma) microenvironment.^13,15,16^ These findings indicate that the inflammatory conditions present in the tumor microenvironment can facilitate the activation of T cells that do not possess T cell receptors specifically targeting cancer antigens. Interestingly, these activated T cells often contribute to tumor cell destruction through alternative cytotoxic pathways, such as interactions involving NKG2D and FAS-FASL.^15–19^ In both viral infections and the tumor microenvironment, activated bystander T cells have been found to express canonical NK cell genes, along with more typical CD8+ effector markers.^14–16,20–22^

Initial evidence for the presence of CD19 CAR-T cell induced bystander activated T cells came from immunohistochemical studies in lymphomas;^13^ however there have thus far been no reports of bystander activation when CAR-T cells are delivered to treat leukemia, a setting where the tumor microenvironment includes the peripheral blood as well as the bone marrow. Furthermore, a comprehensive analysis of the transcriptional profile and the underlying mechanisms governing the activation of bystander T cells during CAR-T cell therapy has remained elusive.

Here, in both non-human primate (NHP) and patient samples, we identify an activated CD8+ CAR^neg^ T cell bystander population characterized by a unique transcriptional profile, including the expression of both T cell activation markers as well as canonical NK cell markers. We further demonstrate that bystander T cells generated *in vitro* can induce significant lysis of leukemic blasts in a TCR-independent manner. These results suggest that bystander T cell activation could contribute to the efficacy of CAR-T cell therapeutics for refractory leukemia.

## Results

### CD20 CAR-T cell expansion results in activation of CD8+ but not CD4+ CAR^neg^ T cells in NHP

To identify CAR^neg^ T cell bystander activation in the peripheral blood after CAR-T cell infusion, we utilized flow cytometry, scRNA-Seq, and single-cell TCR-Seq (scTCR-Seq) to determine whether these cells expanded in the NHP CD20-CAR-T cell model.^23^ As previously described,^23^ this model recapitulates the efficacy and toxicity of human CAR-T cells, including CAR-T cell expansion, induction of B cell aplasia, CRS and ICANS. This model presents a significant advantage, as it consistently demonstrates expansion of CAR-T cells, along with the induction of both clinical and laboratory cytokine release syndrome (CRS), within an immunologically analogous animal system.^23^ Importantly, the activation state of T cells in this model remains unaffected by disease status or prior chemotherapy treatments, enabling a highly uniform platform for transcriptomic analysis.

**Figure 1A** demonstrates the experimental strategy used for a representative NHP, R.315, for which detailed sampling to measure bystander T cells was performed. This included baseline samples (representing sorted T cells collected prior to lymphodepletion and CAR-T cell infusion, termed ‘Pre-infusion T cells’), cells sampled from the infusion product, and cells from the peripheral blood sampled at two time-points (Day 10 and Day 14) after CAR-T cell infusion. The two post-infusion time-points collected included the time of peak CAR-T cell expansion (Day 10) and the time of the start of CAR-T cell contraction (Day 14). For these samples, CAR^pos^ and CAR^neg^ T cells were purified flow cytometrically, identifying CAR^pos^ cells using the CD20-CAR directed Rituximab antibody. **Figure 1B** demonstrates both CAR-T cell expansion/contraction and B cell aplasia in R.315. As shown in the figure, the peak expansion of CAR-T cells in R.315 was 39% of all CD3+ cells at Day 10. This was consistent with CAR-T cell expansion in previously reported NHP CAR-T cell recipients (R.301-304, for which 20%-88% of total CD3+ T cells were CAR+ at peak expansion, **Table S1**).^23^

**Figure 1.**
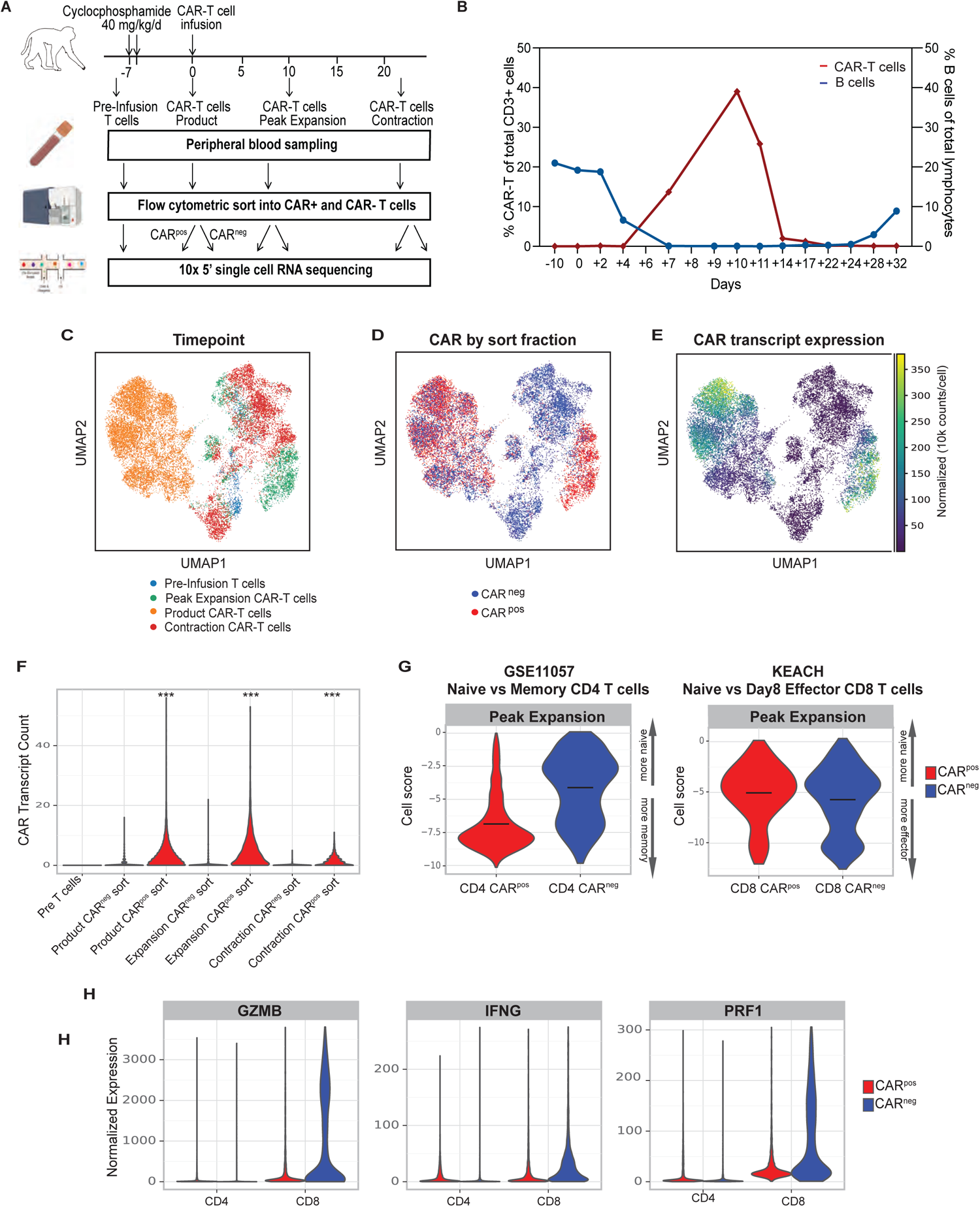
NHP model of CAR-T cell therapy reveals CD8+ CAR^neg^ T cells with an activation signature. (**A**) Schematic of sample preparation: T cells and CAR^pos^ and CAR^neg^ cells were flow cytometrically sorted prior to infusion, from the product, at day 10 (peak expansion), and at day 14 (the start of CAR-T contraction), and prepared for scRNA-Seq and scTCR-Seq. (**B**) CAR-T cell expansion and B cell aplasia were tracked in animal R.315. (**C**) UMAP with shared nearest neighbor clustering of the scRNA-Seq dataset, colored by timepoint. (**D**) UMAP of the scRNA-Seq dataset, colored by flow cytometrically sorted CAR-positive and CAR-negative cells. (**E**) UMAP of the scRNA-Seq dataset, colored by normalized CAR transcript counts. (**F**) Unadjusted CAR transcript counts in sorted CAR^pos^ and CAR^neg^ populations. (**G**) DecoupleR Weighted Sum (WSUM) analysis of a naive vs memory gene signature in CD4 CAR^pos^ and CAR^neg^ T cells, and a naive vs effector gene signature in CD8+ CAR^pos^ and CAR^neg^ T cells, with the analysis performed with cells isolated at the time of peak expansion. (**H**) Violin plots of normalized expression of the CD8+ effector molecules Granzyme B (GZMB), Interferon gamma (IFNG) and perforin 1 (PRF1) in CD4 and CD8+ CAR^pos^ and CAR^neg^ T cells.

To further characterize the CAR^pos^ and CAR^neg^ cells emerging after CAR-T cell infusion, we performed 5’ scRNA-Seq and scTCR-Seq and obtained 11,392 CAR^neg^ and 9,340 CAR^pos^ T cells (after data filtering and quality control) from R.315 (**Figure 1C-F**, **Supplemental Figure 1**). **Figure 1C** demonstrates the separation between the CAR-T cell product and the pre- and post-infusion timepoints comprising both CAR^pos^ and CAR^neg^ T cells purified from the peripheral blood from R.315 via a UMAP plot. **Figure 1D** displays CAR^pos^ (red) and CAR^neg^ (blue) T cells, based on flow cytometrically sorted populations.^23^ **Figure 1E** displays normalized expression of the CAR transcript, with a comparison of **Figure 1D to 1E** confirming high congruency between flow cytometric and transcriptomic identification of the CAR-expressing T cells. **Figure 1F** shows the unadjusted CAR transcript count in each sorted population, with significantly higher levels of the CAR transcript in the CAR^pos^ cells vs CAR^neg^ (Wilcoxon rank sum test, p <0.001), again confirming our ability to identify both CAR^pos^ and CAR^neg^ T cells with high accuracy.

Prior studies have indicated that bystander activation primarily occurs in CD8+ T cells^14–16^. To investigate whether bystander activation during NHP CAR-T cell expansion primarily affected CD8+ T cells rather than CD4+ T cells, we conducted an initial global transcriptional analysis to determine the memory/effector status of CAR^pos^ and CAR^neg^ cells (for both CD4+ and CD8+ T cells) at peak expansion. To accomplish this, we used the DecoupleR^24^ computational package to evaluate the extent to which CD4+ T cells acquired a canonical memory-associated T cell signature (using the GSE11057 gene set^25^), and the extent to which CD8+ T cells acquired a canonical effector-associated T cell signature (using the KEACH Naive vs Day8 Effector CD8+ T cell gene set^26^). **Figure 1G** demonstrates that, for CD4+ T cells, when comparing the peak expansion enrichment scores, CAR^pos^ cells acquired a more memory-like signature at peak expansion compared to CAR^neg^ T cells (mean signature scores at peak expansion: CAR^pos^: −6.86, CAR^neg^: —4.11, p <0.001 using Welch’s t-test for the comparison of CAR^pos^ to CAR^neg^ cells). In contrast, CAR^neg^ CD8+ T cells acquired a similar effector-like signature compared to CAR^pos^ CD8+ T cells (**Figure 1G**), suggestive of activation of bystander CD8+ T cells. Consistent with this effector signature, both CD8+ CAR^pos^ and CAR^neg^ cells demonstrated enrichment for the genes encoding Granzyme B (*GZMB*), Perforin (*PRF1*) and IFNy (*IFNG*) at the time of peak CAR-T cell expansion (**Figure 1H**). These findings indicate that bystander activation predominantly occurred within the CD8+ CAR^neg^ T cell populations.

### Analysis of scRNA-Seq data identifies the transcriptional program of bystander activated CD8+ T cells after CAR-T cell infusion

While alterations in the expression levels of single bystander-defining proteins have been previously characterized ^14–16^, there has not yet been a comprehensive transcriptional analysis to elucidate the underlying mechanisms of activation of these cells. Therefore, to deepen our understanding of the transcriptional signature of NHP CD8+ CAR^neg^ bystander T cells, we performed additional scRNA-Seq analysis on R.315. **Figure 2A** demonstrates Leiden clustering of both CAR^pos^ and CAR^neg^ T cells from this experiment, in which 16 unique cell clusters were identified. Four of these clusters were enriched for CD8+ CAR^neg^ T cells (cluster #s 4, 5,10, 14, **Figure 2B-C**). As shown in **Figure 2C and Supplemental Table S2,** these clusters demonstrated expression of selected effector molecules (*IFNG*, *IL2*), cytolytic molecules (*GZMA, GZMB, GZMM*) as well as several canonical NK cell markers (including *NKG2D* and *KLRD1*). These effector- and NK-associated molecules are similar to those described in bystander activated T cells in the setting of viral infections and in the solid tumor microenvironment.^14–16,20,21^

**Figure 2.**
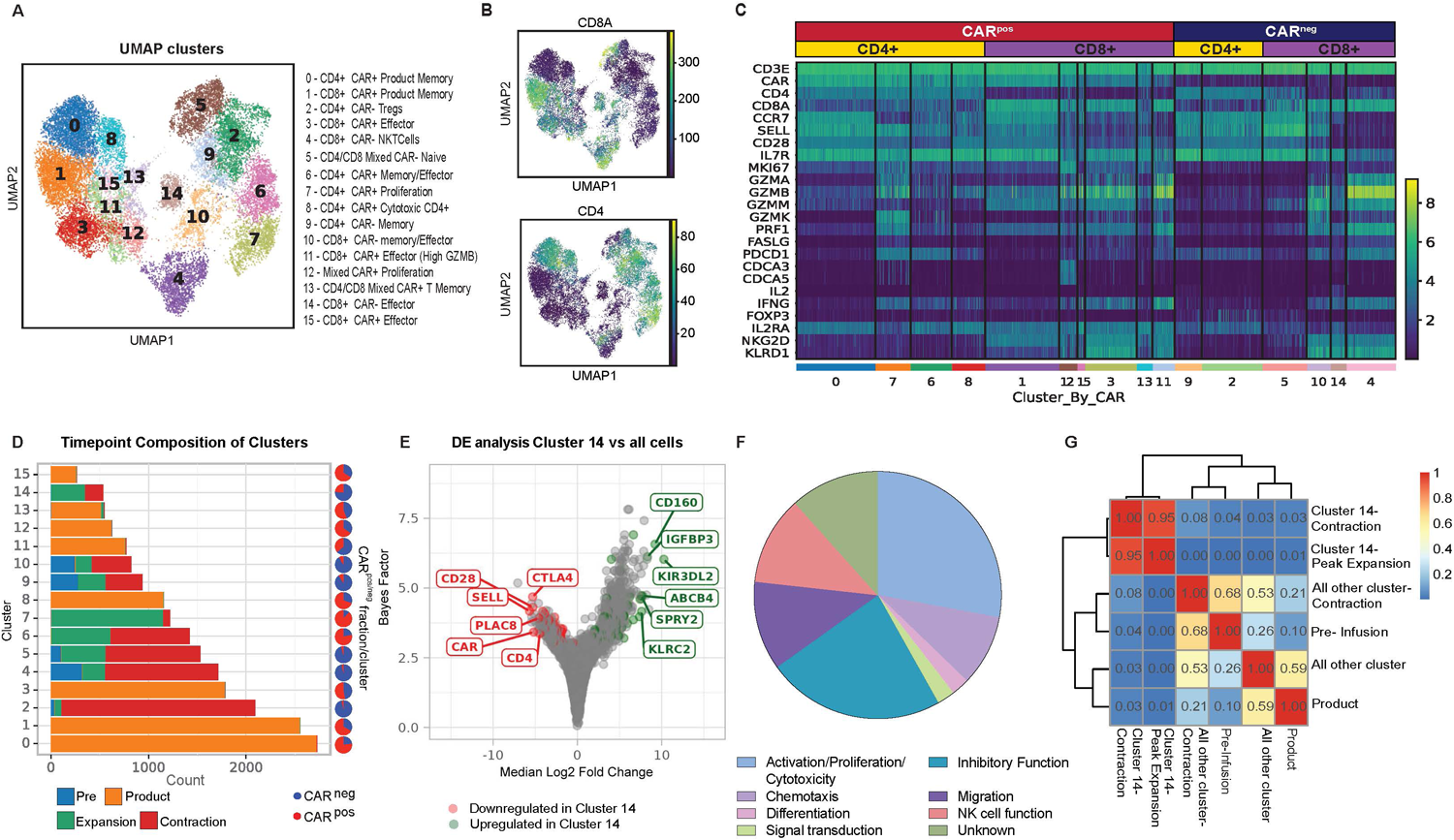
Identification of the CD8+ bystander signature in the NHP model. (A) UMAP with shared nearest neighbor clustering from NHP recipient R.315. 16 Clusters are denoted by colors and labeled with inferred cell states. (B) UMAP with normalized CD8a and CD4 expression. (C) Heatmap demonstrating normalized and log-scaled expression of selected T cell activation, effector and known bystander genes. (D) Cluster composition based on the timepoint of sample collection. (E) Differential expression of the bystander cluster (Cluster 14) vs all other clusters in the dataset. The top six upregulated (green) and downregulated (red) genes by median log2 fold change are labeled. (F) The 42 genes that exhibited upregulation in **Bystander Signature #1** were classified based on their proposed function in T cells. (G) Heatmap of Morisita Index for all samples, comparing cluster 14 cells (at contraction and peak expansion) to all other cells by using the clonotype ID inferred by cellranger to group cells into clones.

To determine which cluster(s) may be enriched in bystander cells associated with CAR-T cell expansion, we investigated the time-point composition of clusters 4, 5, 10, and 14. **Figure 2D** and **Supplementary Table S3** demonstrate that clusters 4, 5, and 10 included a substantial proportion of cells obtained at the pre-infusion timepoint, and therefore were not exclusively identified during CAR-T cell expansion. In contrast, the CAR^neg^ CD8+ T cell Cluster #14 was almost entirely composed of T cells identified post-infusion, suggesting that the effector/activation status of cells in this cluster may be more closely linked to CAR-T cell expansion. Cluster 14 also demonstrated high T Cell Receptor (TCR) diversity (**Supplemental Figure 2A, B)**, consistent with these cells being composed of polyclonal T cells. Further computational analysis therefore focused on Cluster 14.

To develop a transcriptional signature for activated CAR^neg^ CD8+ T cells, we used scVI’s differential expression test.^27,28^ We applied several stringent filters to the results (see **Methods**) to identify a distinct transcriptional signature (‘**Bystander Signature #1’**) of genes that were differentially expressed (DE) in Cluster 14 versus all other cells (see **Methods**). Applying an FDR cutoff of 0.05, this resulted in a list of 43 genes upregulated and 29 genes downregulated in these activated CAR^neg^ CD8+ T cells (**Table 1**). **Figure 2E**, labeled with the top six up- and downregulated genes, demonstrates that this signature included upregulation of T cell effector and NK cell-associated genes, including *CD160* and *KIR3DL2,* and downregulation of naive T cell-associated genes, including *SELL*, *CD28,* and *CCR7* (**Table 1**, **Figure 2E**). This is consistent with a protein expression pattern previously associated with highly activated bystander CD8+ T cells during viral infections and in the tumor microenvironment.^14–16,20–22^ Indeed, an analysis of the 43 upregulated DE genes underscored this conclusion, with 40% of these genes being associated with effector T cell and/or NK cell function (**Figure 2F**). The differential expression test used for **Bystander Signature #1** measured overall differences between all bystander CD8+ T cells and all non-bystanderT cells (both CD4+ and CD8+) in our dataset. To more specifically identify what distinguished bystander CD8+ T cells from other CD8+ T cells, we conducted additional bystander vs non-bystander DE tests: We performed DE analysis only on CD8+ T cells (creating the **‘Bystander Signature #2’, Supplemental Figure 2C**, **Supplemental Table S4**), and only on CAR^neg^ CD8+ t cells at the time of peak CAR-T cell expansion (creating **‘Bystander Signature #3’, Supplemental Figure 2C**, **Supplemental Table S4**). All three signatures demonstrated substantial overlap of upregulated and downregulated genes (**Supplemental Figure 2D**), underscoring the consistency of these analyses in identifying bystander-defining genes. Finally, we created a minimal gene list, **‘Bystander Signature #4’**, that enabled the identification of the bystander cluster (Cluster 14, (**Supplemental Figure 2C**). This signature was generated using five genes (*CD8A*, *CD160*, *KLRK1* (NKG2D), *KLRD1* (CD94) and *CCL5*) that are each highly expressed in the bystander cluster and that are also amenable to flow cytometric identification. Underscoring the rigor of our cluster identification, the CAR^neg^ bystander cluster was also distinct across multiple levels of clustering resolution. Thus, the same cluster of bystander cells was identified when the data were clustered using the Leiden algorithm^29^ at resolutions of 0.8,1.0 and 1.2 (**Supplemental Figure 2E, F**).

**Table 1:** Gene list for Bystander Signature #1. Listed genes are differentially expressed (up and downregulated) in Cluster 14 in comparison to all other cells in the dataset for animal R.315.

| Upregulated genes |  | Downregulated genes |  |
| --- | --- | --- | --- |
| gene name (human ortholog) | log fold change | gene name | log fold change |
| *KIR3DL2* | 10.26 | *EZH2* | -1.73 |
| *CD160* | 9.21 | *SLC2A3* | -1.78 |
| *IGFBP3* | 8.31 | *GNG2* | -1.83 |
| *ABCB4* | 7.72 | *LYST* | -1.97 |
| *ENSMMUG00000050862 (KLRC2)* | 7.62 | *LTB* | -1.98 |
| *SPRY2* | 7.58 | *TESPA1* | -1.98 |
| *XCL1* | 7.44 | *SATB1* | -2.11 |
| *CCL5* | 7.36 | *CCR7* | -2.16 |
| *NUGGC* | 6.65 | *ICOS* | -2.24 |
| *ENSMMUG00000063583 (GNLY)* | 6.63 | *SIT1* | -2.25 |
| *ENSMMUG00000013779* | 5.95 | *DUSP16* | -2.38 |
| *CCR9* | 5.93 | *SH2D1A* | -2.41 |
| *KLRD1* | 5.52 | *PTPRJ* | -2.43 |
| *ENSMMUG00000054501 9 (TRGV)* | 5.47 | *H1-3* | -2.52 |
| *CD101* | 5.04 | *SLFN12L* | -2.71 |
| *PPP2R2B* | 4.89 | *PDE4B* | -2.71 |
| *SERPINB6* | 4.61 | *PGAP1* | -2.72 |
| *PLCG2* | 4.55 | *CFP* | -2.75 |
| *ENSMMUG00000057791 (TRDV)* | 4.54 | *ANTXR2* | -2.88 |
| *LAT2* | 4.48 | *GPR183* | -2.93 |
| *ANK3* | 4.16 | *SPOCK2* | -3.37 |
| *THAP6* | 4.03 | *MT1E* | -3.46 |
| *JAKMIP2* | 4.02 | *S1PR1* | -3.89 |
| *THY1* | 3.96 | *LEF1* | -4.09 |
| *MATK* | 3.87 | *TOB1* | -4.13 |
| *ENTPD1* | 3.71 | *CD4* | -4.47 |
| *ENSMMUG00000032338 (SMIM10)* | 3.62 | *PLAC8* | -4.59 |
| *TIGIT* | 3.59 | *CD28* | -5.29 |
| *OTUD5* | 3.52 | *SELL* | -5.68 |
| *FCRL6* | 3.44 |  |  |
| *NCALD* | 3.32 |  |  |
| *NKG7* | 3.17 |  |  |
| *ITGA1* | 3.03 |  |  |
| *NR3C2* | 2.88 |  |  |
| *DAPK2* | 2.79 |  |  |
| *SPINK2* | 2.73 |  |  |
| *PRR29* | 2.73 |  |  |
| *STX3* | 2.34 |  |  |
| *RIN3* | 2.32 |  |  |
| *NR4A1* | 2.27 |  |  |
| *S100A6* | 2.23 |  |  |
| *UBASH3B* | 2.13 |  |  |
| *CPQ* | 2.11 |  |  |

### Bystander T cells arise from recipient peripheral blood CD8+ T cells after CAR-T cell infusion

To determine whether CD8+ CAR^neg^ bystander T cells expanded from the infused product, or from T cells in the recipient, we performed VDJ-TCR single cell sequencing and assembled TCRs^30^ from R.315, assigned them to clones, assessed their number and diversity, and determined their similarity across samples (**Supplemental Figure 2 A,B**). We utilized the Morisita index, an ecological metric commonly used to assess overlap of species counts between two different environments^31^, to examine the degree of clonal overlap between Cluster 14 clones and clones identified in the infused product, at the time of peak CAR-T cell expansion, and in the CAR-T cell contraction phase. This analysis identified a high degree of clonal overlap between Cluster 14 CAR^neg^ bystander T cells present at the time of CAR-T peak expansion and contraction, but much less overlap between this cluster and either the pre-infusion T cells or the infused product (**Figure 2G**). These data support a model wherein the majority of the CAR^neg^ bystander cells arise from peripheral blood CD8+ T cells that acquire this unique phenotype in the setting of CAR-T cell expansion.

### Bystander T cells identified in 4 additional NHP recipients of CD20-CAR T cells

To determine if cells bearing the bystander signature were present in other NHP recipients of CAR-T cells, we applied **Bystander Signature #1** to a validation cohort of four additional recipients.^23^ The clinical and CAR-T course for these animals has been previously described,^23^ with CAR-T cell expansion in these animals being comparable to animal R.315 (**Supplemental Table 1**). For this analysis, we again performed 5’ scRNA-Seq on pre-infusion T cells, CAR^pos^ and CAR^neg^ T cells from the infused CAR-T product, from the peripheral blood at the time of peak CAR-T expansion, and from the peripheral blood at the onset of CAR-T contraction **(Supplemental Figure 3 A-C)**. **Figure 3A** demonstrates the Leiden clustering of 29,050 CAR^neg^ and 5,480 CAR^pos^ derived from animals R.301, R.302, R.303 and R.304 with 24 distinct clusters identified (**Supplemental Tables S5 and S6**).

**Figure 3.**
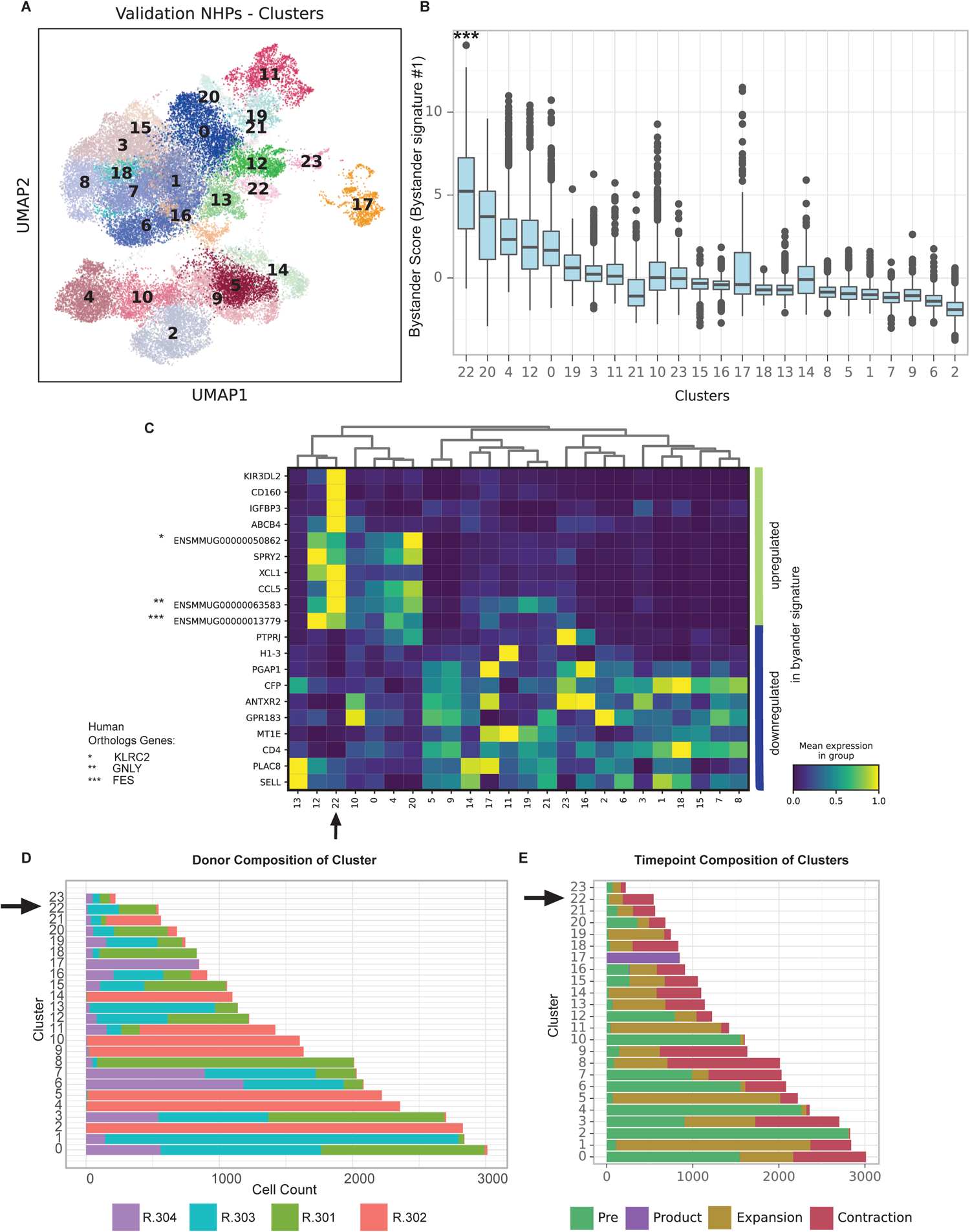
Bystander CAR-T cells can be detected in a validation cohort of 4 additional NHP. (A) UMAP with shared nearest neighbor clustering across 4 additional animals, colored to identify 23 transcriptional clusters. (B) Bystander score determined by applying **Bystander Signature #1** to each cluster using DecoupleR. (C) Heatmap of log-scaled normalized expression, averaged across each cluster, for the top ten upregulated and downregulated genes identified in **Bystander Signature #1**. (D) Composition of cluster by animal, with an arrow highlighting Cluster #22, the cluster with the highest signature score when applying **Bystander Signature #1.** (E) Composition of clusters based on sample collection timepoint, with an arrow highlighting Cluster #22, the cluster with the highest signature score when applying **Bystander Signature #1**.

We next applied **Bystander Signature #1** to these clusters, using DecoupleR to score the cells. As shown in **Figure 4B**, Cluster 22 exhibited a much higher **Bystander Signature #1** score than the other clusters, identifying the cluster as likely consisting of bystander T cells (Cluster 22 signature score vs score on all other cells, Welch’s t-test p<0.001). Cluster 22 also achieved the highest signature scores when applying **Bystander Signatures # 2-4** (**Supplemental Figure 3D**, Welch’s t-test p<0.001 for all three signatures). The **Figure 3C** heatmap displays the log-scaled normalized expression for the top ten upregulated and bottom ten downregulated genes for **Bystander Signature #1**, illustrating the alignment of Cluster 22 with this signature. A plot of all available genes in the dataset shared with the signature (**Supplemental Figure 4A**) shows similar concordance. **Figure 3D** demonstrates that Cluster 22 is composed of cells from all four animals albeit with two animals dominating the cluster (R.303, R301), and with the majority of cells (95%) coming from the maximal expansion and contraction timepoints (**Figure 3E**), consistent with the findings from R.315 (**Figure 2**).

**Figure 4.**
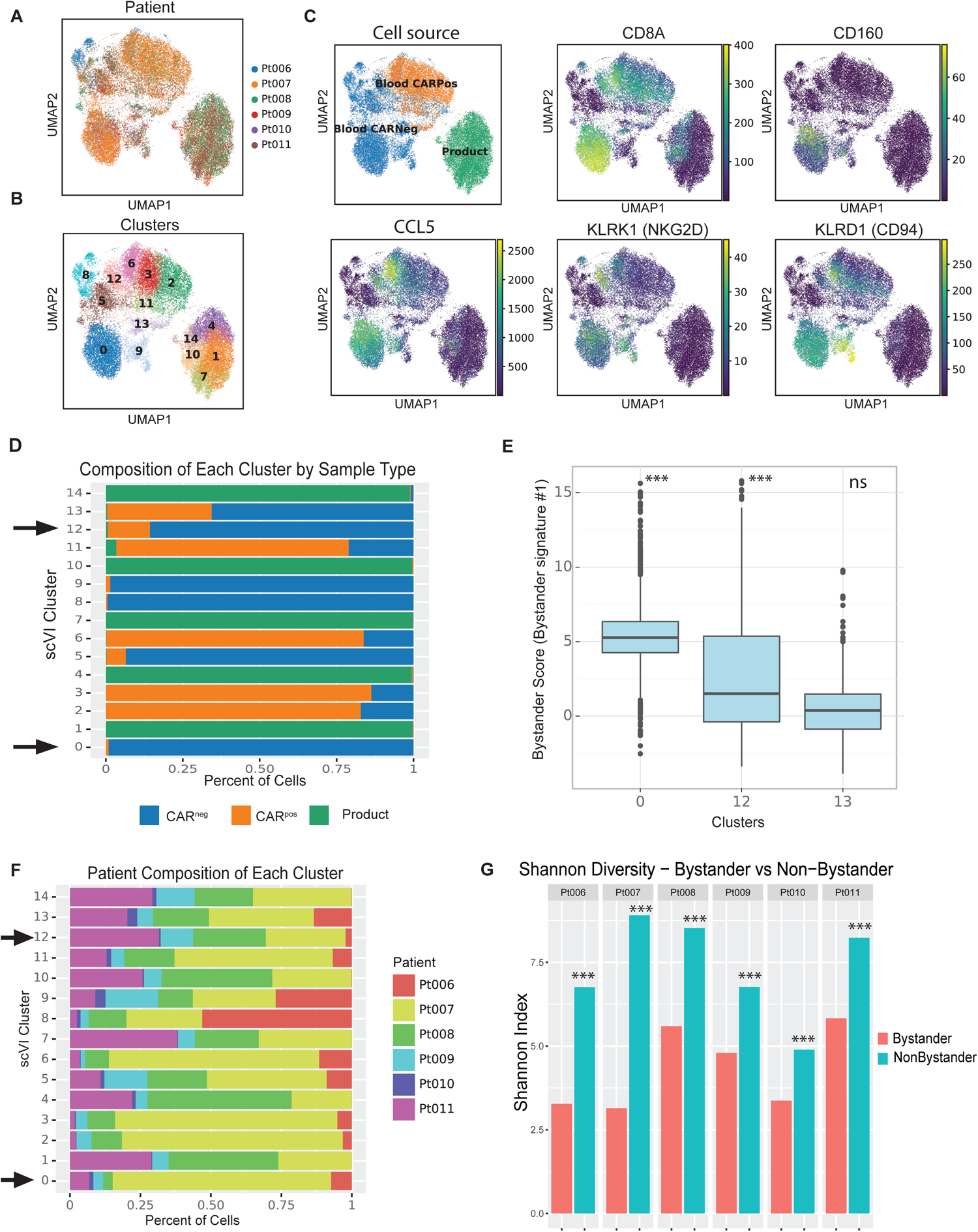
Identification of CD8+ CARneg bystander T cells in patients after Tisangelecleucel infusion. (A) UMAP of scRNA-seq data from six pediatric patients post Tisangelecleucel, colored by patient ID. (B) UMAP with shared nearest neighbor clustering across all patients, colored by 15 transcriptional clusters. (C) UMAP colored by cell source, as well as normalized expression of genes from **Bystander Signature #4** (CD8+, *CD160, CCL5, KLRK1* (NKG2D) and *KLRD1* (CD94). The product was not cytometrically sorted into CAR^pos^ and CAR^neg^ T cells and therefore contained a mixed population of CAR^pos^ and CAR^neg^ T cells. (D) Composition of clusters by cell source, with arrow highlighting Clusters 0 and 12, which were enriched for **Bystander Signature #1**. (E) Bystander score determined by applying **Bystander Signature #1** to each CD8+ CAR^neg^ cluster using DecoupleR. (F) Composition of clusters by Patient ID, with arrows highlighting Clusters 0 and 12, which were enriched for **Bystander Signature #1**. (G) Shannon diversity of T cell clones in Bystander clusters 12 and 0 vs all CD8+ CAR^neg^ cells in each patient.

### Bystander activated T cells are present in patients after Tisangelecleucel infusion

To determine if CD8+ CAR^neg^ bystander T cells were also activated in patients receiving CAR-Ts for acute B cell lymphoblastic leukemias (B-ALL), we utilized samples from six pediatric patients with a diagnosis of relapsed B-ALL who received Tisangelecleucel, and who were enrolled on an institutional biology study (**Supplemental Table S7**). We obtained 11,555 total T cells from the CAR-T products infused into these six patients, and 23,322 T cells from peripheral blood collected on Day +6 after infusion. The peripheral blood cells were sorted into CAR^pos^ T cells (9,073 total cells) and CAR^neg^ T cells (14,249 total cells) (**Supplemental Figure 5 A-C**). We then performed 5’ scRNA-Seq and Leiden clustering, which identified 15 distinct clusters (**Figure 4B**, **Supplemental Tables S8 and S9**).

To identify potential bystander T cell clusters, we first identified CD8+ clusters composed of more than 50% CAR^neg^ T cells (Clusters 0, 12, and 13) and then scored these cells using **Bystander Signature #1**. We thereby identified three potential clusters (Clusters 0,12, and 13, **Figure 4B-C**, **Supplemental Figure 5D**), each of which was primarily composed of CAR^neg^ T cells collected after CAR-T infusion, with each cluster being composed of <1% T cell clones from the infused cell product (**Figure 4D**). The mean score for **Bystander Signature #1** on these clusters were Cluster 0: 5.37, Cluster 12: 2.82, and Cluster 13: 0.44. Applying Welch corrected t-tests to compare whether the mean **Bystander Signature #1** score was greater in each cluster compared to all of the remaining cells, we found that Clusters 0 and 12 each were significantly enriched (p <0.001) for **Bystander Signature #1**, but that Cluster 13 was not (p=1.0, **Figure 4D**). Of note, the analysis depicted in **Figure 4** included Patient ID as a batch term to account for patient specific clusters. As shown in **Figure 4F**, Cluster 0 was predominantly composed of cells from Patient #Pt-007 (77.5%) while 85.7% of Cluster 12 cells included cells from patients #Pt-007, #Pt-008, and #Pt-011. Repeat analysis was also conducted without including individual patient IDs as a batch term, and yielded similar results (**Supplemental Figure 6**, **Supplemental Tables S10-S11**). Thus, as shown in **Tables S12-S13**, the cells identified as bystanders were quite similar in both analyses, with a Jaccard Index=0.72 for the two datasets (**Table S12**) and the share of T cells identified as bystanders similar for most patients in the dataset (**Table S13**). Of note, Shannon diversity analysis of the CD8+ CAR^neg^ bystander cells (Cluster 0 and 12) vs non bystander cells (Cluster 13) demonstrated a lower clonal diversity in the bystander clusters within each patient (Hutcheson’s t-test within each patient, all p-values < 0.001) (**Figure 4G**), which was driven by expansion of several large clones (**Supplemental Table S14 and S15**). While some CDR3 regions of these large clones were commensurate with published CMV or EBV specific TCRs ^32,33^, the majority of the clones in this cluster were uncharacterized. Therefore, the antigen-specificity of these expansions remains to be determined.

### Bystander activation is induced by the gamma cytokines IL-2 and IL-15

The identification of CD8+ bystander CAR^neg^ T cells in both NHP and patients prompted us to explore the mechanisms that might contribute to their activation. In viral infection models, stimulation of bystander T cells with cytokines has been proposed as the major activating mechanism.^14^ To test this hypothesis *in vitro*, we exposed healthy donor human T cells to cytokines that have previously been documented to be elevated after CAR-T infusion^34,35^. These included IL-6, IFN-y, GM-CSF, IL-2, IL-15, IL-7, IL-18, IL-12, IL-4, IL-10 and TNF-a. We determined whether exposure to these cytokines would increase expression of some of the markers identified on the bystander CAR^neg^ CD8+ T cells, including CD8, CD160, NKG2D, and CCL5 (**Figure 5A**). **Figure 5B** demonstrates that only stimulation with the gamma cytokines IL-2 and IL-15 was able to increase the expression of CD160, NKG2D, and CCL5 in CD8+ T cells, consistent with the acquisition of the bystander CD8+ phenotype. Of note, TCR stimulation with CD3/CD28-coated beads did not result in the evolution of CD8+ T cells towards a bystander phenotype, suggesting that direct TCR stimulation is not responsible for the generation of this cell population. Titration of IL-2 (0-1000 U/ml) and IL-15 (0—50ng/ml) demonstrated a dose­dependency of the expression of the bystander markers CD8, CD160, NKG2D, and CCL5. Addition of 500U/ml IL-2 and 50ng/mg IL-15 resulted in the largest population of CD8+ T cells with expression of the bystander markers CD8, CD160, CCL5 and NKG2D (15.5% +/− 6.6, 16.1% +/−5.4, respectively) when compared to all other conditions (**Figure 5C and D**).

**Figure 5.**
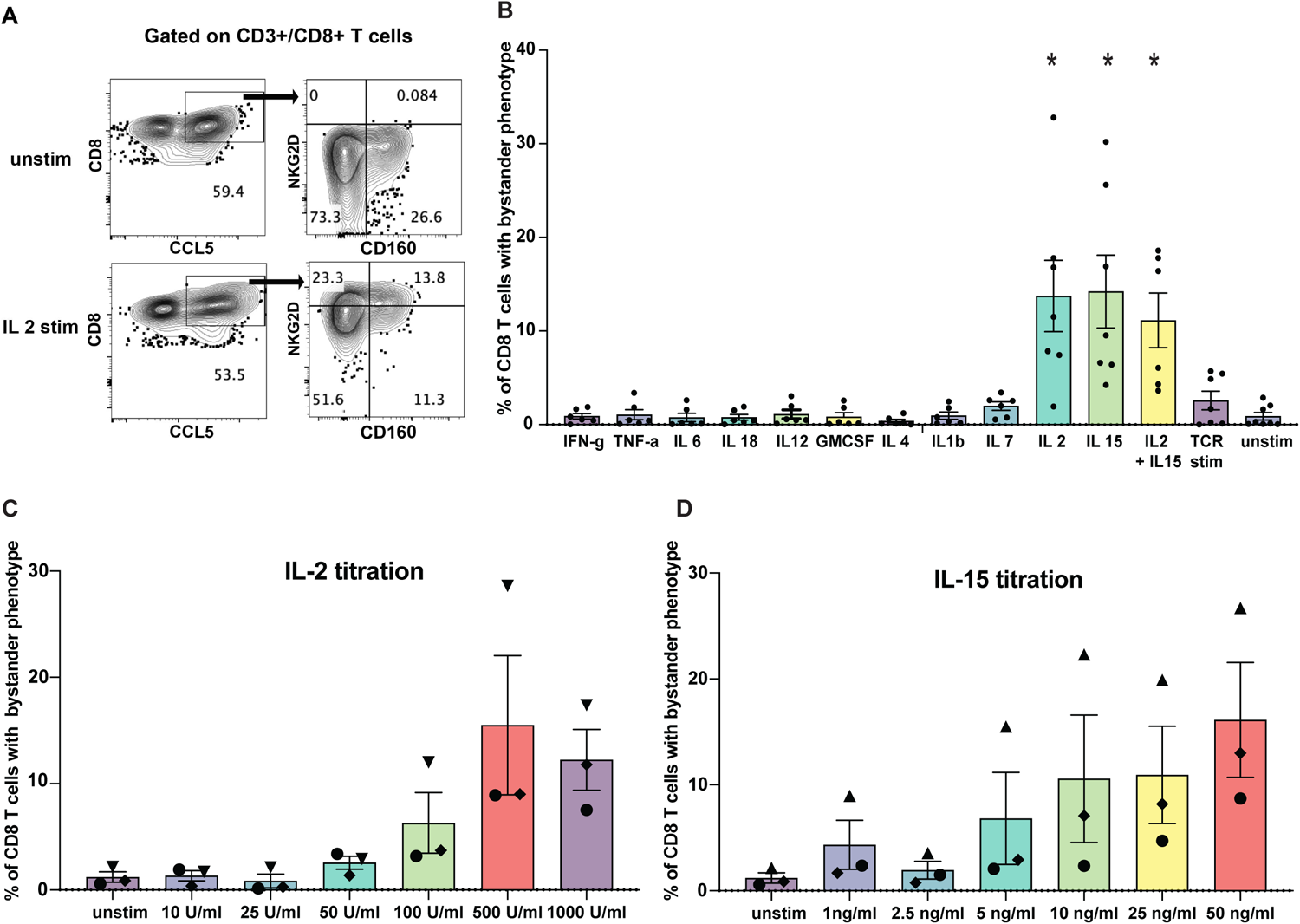
Stimulation of primary human T cells with the gamma cytokines IL-2 or IL-15 generates cells with a bystander phenotype. (A) Representative flow plots demonstrating the gating strategy used to identify T cells expressing the bystander markers CD8+, CD160, NKG2D, and CCL5 in unstimulated cells (unstim), and cells stimulated with IL-2. Cells were gated on CD3+/CD8+ double positive T cells and then subsequently gated on the bystander markers CCL5, NKG2D and CD160. CD8/CCL5 double-positive cells were assessed for the expression of the bystander marker of NKG2D and CD160. (B) Percentage of total CD8+ T cells exhibiting expression of the bystander markers CD160, NKG2D and CCL5 after stimulation with CRS-associated cytokines. (C) Percentage of total CD8+ T cells with the bystander phenotype after titration of IL-2 and (D) IL-15.

To assess if bystander CD8+ T cells could potentially participate in tumor control, healthy donor primary human CD8+ T cells were activated with IL-15, and then flow-sorted based on the presence or absence of the expression of the bystander markers NKG2D, CD94 and CD160 (**Figure 6A**). NKG2D/CD94/CD160-positive and -negative T cells were then co-cultured with the B-ALL leukemia cell line Nalm6 (**Figure 6B**). Evaluation of live Nalm6 cells by flow cytometry 16 hours after co-culture revealed a significantly higher degree of tumor killing by bystander NKG2D/CD94/CD160-positive CD8+ T cells compared to NKG2D/CD94/CD160-negative T cells (80.96% killing +/− 5.7 vs 58.2 % killing +/− 9.7%, p<0.01, paired t-test, **Figure 6B**). To assess if this killing was TCR-dependent or -independent, sorted bystander CD8+ T cells were also cocultured with p2M knock-out Nalm6 cells (lacking MHC-I surface expression). Similar amounts of target cell lysis were observed when these cells were co-cultured with bystander CD8+ T cells (98.2% +/− 0.7% killing by NKG2D/CD94/CD160-positive cells vs 63.9%+/− 1.9% by NKG2D/CD94/CD160-negative cells, p<0.01, paired T-test), consistent with a TCR-independent cytotoxic mechanism (**Figure 6C**).

**Figure 6.**
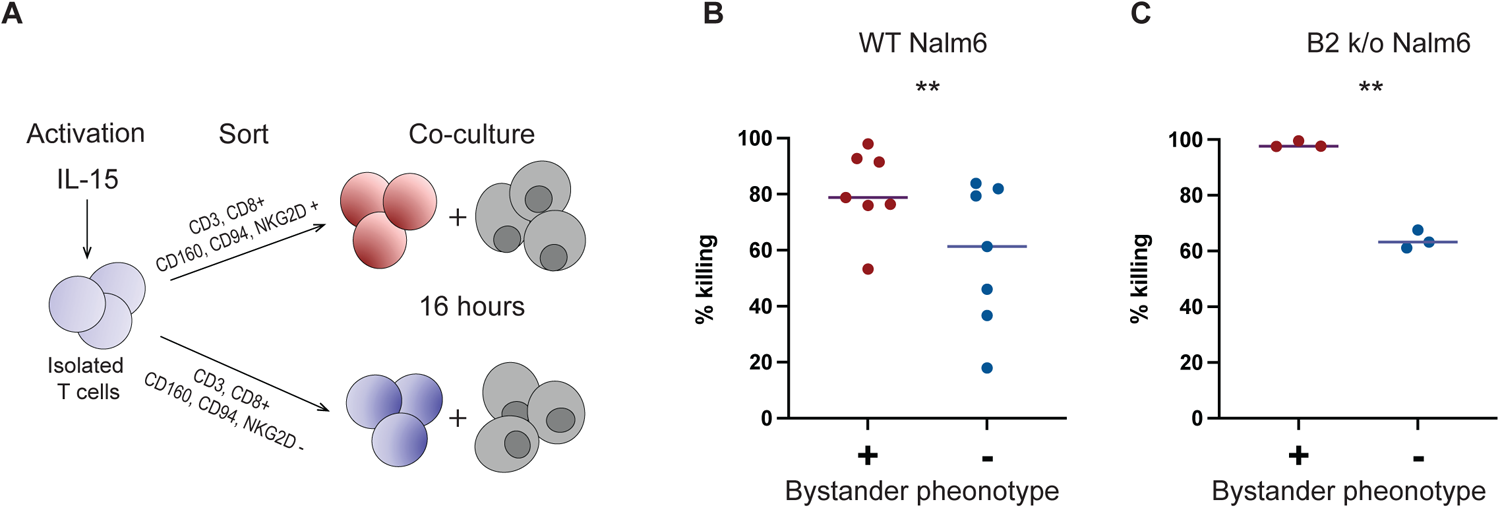
Cytokine-stimulated primary human T cells expressing bystander surface markers are capable of killing leukemic cells. (A) Schematic of co-culture cytotoxicity assay (B) Percentage killing of Nalm6 and (C) Nalm6 b2M k/o cells 16 hours after co-culture with sorted CD8+ T cells, with and without the bystander phenotype at a target:effector ratio of 1:1. Percentage killing was calculated by comparing the remaining live NALM6 cells in cultures containing CD8+ T cells to the remaining live Nalm6 cells in cultures without the addition of T cells.

## Discussion

Activation of bystander T cells has been identified in the setting of viral infection, as well as in the solid tumor microenvironment.^14–16,36^ However, bystander activation in the setting of CAR-T cell therapy for leukemia has not yet been described. Here we demonstrate that B cell-directed CAR­T cell infusion results in bystander activation of CD8+ T cells in NHP, and in patients receiving Tisangelecleucel. We further identified bystander-specific gene signatures which were enriched for genes involved in T cell activation, in CD8+ effector functionality, and which included several canonical NK cell-associated genes. Furthermore, we demonstrated that bystander activation could be induced by IL-2 and IL-15, and that bystander activated cells have the potential to kill leukemic cells in a TCR-independent manner.

T cell bystander activation has been studied in viral infections including hepatitis B,^37,38^ Influenza^39^, CMV^21^ and HIV.^40,41^ In these settings, bystander activated, non-viral antigen specific T cells were found to be CD8+ effector T cells which also demonstrated expression of NK cell markers including NKG2D, CD56, KIR, NKG2A, NKG2C, CD94, NKp30, expression of effector cytokines including IFN-y, as well as cytolytic enzymes, including Granzyme B and Perforin. ^13–16,20,21,36,39,42^ Bystander activated T cells of a similar phenotype have also been identified in multiple solid tumors, including the identification of activated CAR^neg^ T cells infiltrating the tumor microenvironment of diffuse large B cell lymphomas in patients who received the CD19 CAR-T cell therapeutic Axicabtagene.^13^ Although multiplex immunohistochemistry assays revealed that these CAR^neg^ T cells were CD8+ T cells expressing proliferation, proteolytic, and activation molecules^13^, there is still a lack of comprehensive phenotypic and transcriptional analysis of activated bystander T cells residing in the peripheral blood, outside of the tumor microenvironment, in the context of CAR-T cell therapy for liquid tumors.

As patient-derived datasets for transcriptional analysis can be influenced by factors including disease stage and treatment history, we first employed a NHP model of CAR-T cell therapy to explore the activation of CAR^neg^ bystander cells. This model provided a consistent framework for identifying transcriptional patterns specific to bystander effects, and enabled the discovery of a series of four Bystander Signatures which subsequently served as the foundation for the analysis of patient samples. The phenotypic markers of bystander activated CD8+ T cells identified in this study demonstrate substantial overlap with bystander CD8+ T cells identified in viral infections and solid tumors.^14–16^ Importantly, in addition to standard phenotypic evaluation, the current analysis included an in-depth single-cell transcriptomic analysis of these cells, which is expected to expand and deepen our ability to understand the broader T cell immune environment that becomes activated during CAR-T therapy.

Bystander-activated T cells have been proposed to enhance both antiviral and antitumor immune responses.^14,15^ This enhancement is believed to be partly mediated through TCR-independent cytotoxicity, involving molecules such as NKG2D or the Fas-FasL interaction.^17,18,20,42,43^ Our experiments also confirmed TCR-independent killing of leukemic cells. However, within the tumor microenvironment, some bystander-activated T cells may possess tumor antigen specificity and engage in TCR-dependent mechanisms of killing. It is notable that in the NHP CAR-T model, the target cells are autologous non-malignant B cells, which are not expected to express leukemia-derived neo-antigens, and therefore are not anticipated to trigger non-CAR-T cell antigen-specific clonal expansion of T cells. Our finding that CAR^neg^ bystander cells in this non-tumor model did not include prominent clonal expansions is consistent with this fact. In contrast, in the patient samples analyzed, we did identify clonal expansions within the bystander population. While some clones were commensurate with published CMV or EBV specific TCRs, the specificity of the majority of these clones could not be determined through public databases. The determination of whether these clones potentially possess anti-leukemic specificity remains an important area for future investigation.

In this study, we identified the gamma cytokines IL-2 and IL-15 as the major drivers of *in vitro* differentiation of CD8+ T cells towards the bystander activated phenotype. This finding suggests that ongoing clinical trials of CAR-T cells that include the administration of IL-15, or the development of CAR-Ts that autologously express this cytokine^44–47^ may have a salutary effect on bystander as well as CAR-T cells. This may be of particular importance given that our *in vitro* data demonstrated the potential for anti-leukemic cytotoxicity of these bystander T cells. Further investigations utilizing large patient datasets to assess the association of bystander expansion with the duration of clinical response to CAR-T cells will be required to validate the ability of these cells to enhance clinical tumor control. Our identification of a bystander signature that is amenable to both transcriptomic and flow cytometric assessment represents an important milestone towards conducting future analyses for this purpose.

In summary, this study presents a comprehensive analysis of CD8+ bystander T cell activation in the setting of CAR-T cell expansion in both NHP and in patients. It reveals a unique gene expression signature of these cells, and their antigen-agnostic anti-tumor capacity. The identification of a robust bystander CD8+ T cell signature establishes a critical foundation for future analyses of these cells in patients undergoing B cell-directed CAR-T cell therapeutics.

## Methods

### CD20 CAR vector and virus production

#### Two different NHP CD20 domains were utilized in this study

Animal R.315 received a CD20 CAR with an antibody domain identical to the clinically-used CD20 antibody Rituximab.^48^ The remaining animals (R.301-304) received a CD20 CAR-T cell product with a previously described CD20 CAR antibody domain.^23^ All CAR-T cell vectors expressed a CD28 transmembrane domain, a 4-1BB costimulatory domain and the CAR construct utilized for animals R.301-304 additionally expressed EGFRt.^23^ CARs were encoded by an xHIV plasmid which was co-transfected with an HIV-1 Rev/Tat and VSV-G envelope plasmid (RRID:Addgene_138479) for lentiviral production as previously described.^23^

### Transduction and Expansion of NHP CD20 CAR-T cells

NHP CD20 CAR-T cells were transduced and expanded as previously described:^23^ Peripheral blood mononuclear cells (PBMC) were isolated from adult NHP peripheral blood by Ficoll-PaquePLUS (GE Healthcare Bio-Sciences) or SepMate-50 (Stemcell). Total T cells were isolated from PBMCs using a T cell isolation kit per manufacturer’s instructions (Miltenyi Biotech). Polyclonal T cells were activated with Miltenyi NHP T cell Activation/Expansion kit (Miltenyi Biotec) in NHP media (X-Vivo 15 medium (Lonza) supplemented with 10% FBS (HyClone/Gibco), and recombinant human IL-2 (rhIL2,50 U/mL; R&D Systems/CellGenix)) with or without 1% penicillin/streptomycin/l-glutamine (Invitrogen), with or without 50 pmol/L p-mercaptoethanol (Sigma). Lentiviral transduction with spinoculation was performed 24-48 hours post stimulation. At the end of the stimulation cycle, stimulation beads were removed using Miltenyi LS columns (Miltenyi Biotec) and cryopreserved.

### NHP In Vivo Adoptive T cell Transfer Studies

NHP experiments were performed according to the Guide for the Care and Use of Laboratory Animals of the National Institutes of Health, which were approved by the University of Washington, Boston Children’s Hospital, and the Biomere Institutional Animal Care and Use Committee. Each Rhesus Macaque recipient received cyclophosphamide (Baxter) as lymphodepleting chemotherapy at a dose of 40 mg/kg/dose cyclophosphamide on two consecutive days (day −7 and −6). Mesna (Sagent Pharmaceuticals) was administered to each recipient as a bladder protectant. The total mesna dose was equal to the cyclophosphamide dose, divided into 4 doses (7.5-10 mg/kg/dose), administered intravenously (i.v.), 30 minutes prior, and 3, 6, and 8 hours after cyclophosphamide infusion. CD20 CAR-T cells were infused at doses ranging from 0.6×10^7^ to 1.2×10^7^ CD20 CAR-T cells/kg. All recipients received antibiotic prophylaxis with ceftazidime and vancomycin, antiviral prophylaxis with acyclovir and weekly cidovovir and antifungal prophylaxis with fluconazole. Recipients underwent clinical and neurologic monitoring according as previously described.^23^ Peripheral blood was collected longitudinally in all recipients and cryopreserved for later use.

### Patient sample collections and cryopreservation

Peripheral blood samples from pediatric patients with B-ALL receiving Trisangeleucel were collected under the Boston Children’s Hospital and Dana Farber Cancer Institute approved clinical protocol, ‘PREDICT’ (NCT03369353). Patient blood was obtained and PBMCs were isolated by Ficoll-PaquePLUS (GE Healthcare Bio-Sciences) gradient centrifugation. PBMCs were cryopreserved and stored for further use.

### Human and NHP T cell flow cytometric sorting

Cryopreserved NHP and human cells were thawed, stained with fixable Live/Dead stain (Invitrogen), CD4 (BD bioscience, Cat# 563914, RRID:AB_2738485), CD8 (BD bioscience Cat# 563795, RRID:AB_2722501), CD20 (BD Bioscience, Cat# 566988, RRID:AB_2869992) and CD14 (Biolegend, Cat# 301834, RRID:AB_11126983) antibodies. The CAR was detected by staining with an anti-rituximab conjugated flow antibody (GeneTex, Cat# GTX43347, RRID:AB_11166924) in animal R.315 and with an anti-EGFR antibody (R&D systems, Cat# FAB9577G-100) for animals R.301-304. CD19 CAR-T cells from patient samples were stained with an anti-CD19 CAR antibody (ACRO Biosystem, Cat#: CD9-HF251-25ug). CD3 flow cytometric antibodies were not utilized, to avoid in vitro stimulation, which can result in alteration of the transcriptional profile. Live CARpos and CARneg, CD4/CD8pos and CD20/CD14neg cells were flow cytometrically sorted and immediately processed for single cell sequencing.

### Library Preparation

All products used in the preparation of these samples, excluding the NHP TCR specific custom primer set^30^, are available from 10x Genomics. Samples in this publication were prepared using the Next GEM 5’ v2 Gel Beads with the i7 Multiplex Plate (Single Index) or the Next GEM Single Cell 5’ v2 Sample Index Plate TT (Dual Index). GEMs were generated using v2 Gel Beads and the v1 Target Enrichment kit utilizing the off-the-shelf human/mouse T Cell Mix 1 and 2 premixed primers for the human samples and substituting the provided primers with custom designed primers as previously described^30^ for NHP samples.

### Cytokine stimulation assays

For the cytokine stimulation assay, healthy donor PBMCs were obtained under a Boston Children’s Hospital IRB-approved healthy donor protocol. PBMCs were selected as previously described and sorted for T cells using the Pan T cell isolation kit (Miltenyi Biotech). T cells were plated at 200,000 cells in 96 well U bottom plates and stimulation with the following cytokines for 24-48 hours: IL-2 (10-1000U/ml), TNF-a (100ng/ml), IL-15 (1-50ng/m)l, IL-10 (100ng/m)l, IL-12 (100ng/ml), IL-6 (1000ng/ml), GM-CSF (8000 U/ml), IL-4 (10000U/ml), IL-7 (250ng/ml), IL-18 (1000ng/ml) and IFN-y (1.25 ug/ml). After cytokine stimulation cells were collected for flow cytometry and stained with CD3, CD4, CD8, CD160, NKG2D, CD94, CCL5.

### Cytotoxicity assays

Healthy donor PBMCs were isolated as described above. Cells were stimulated with IL-15 (50 ng/ml) for 24 hours. After 24 hours, cells were stained with a Live/Dead marker, CD3, CD4, CD8, CD94, NKG2D, CD160. CD8+ cells were sorted on the live CD3, CD8, NKG2D, CD160 and CD94 positive or negative populations. Sorted cells were counted and placed in coculture with Nalm 6 (Nalm6 Clone G5, ATCC Cat# CRL-3273) and Nalm 6 b2M k/o T cells at an effector: target ratio of 1:1. After 16 hours, Precision Count Beads (Biolegend, Cat# 424902) were added at a 1:1 ratio to target cell input. Cells were collected and stained with Live/Dead, Caspase3+7, CD3 and CD19. The percentage killing was determined by the number of live, Caspase 3+7 negative, CD19+ Nalm6 cells.

### scRNA-Seq analysis

Details of scRNA-Seq analysis are found in **Supplemental Methods**.

