## Supplemental Material for "B cell directed CAR-T cell therapy results in activation of CD8+ cytotoxic CAR-negative bystander T cells in both non-human primates and patients"

**Supplemental Methods:**

**scRNA-Seq analysis:**

**GEX Alignment and TCR Reconstruction**

Paired GEX and VDJ libraries were aligned using the 'Cellranger multi' pipeline (v7.0.0) with 32 CPU's and 128 GB of RAM, with VDJ libraries listed as "vdj-t" to indicate T cell libraries. Samples were aggregated with the 'Cellranger aggr' pipeline (v7.0.0) with arguments '--normalize=none --nosecondary'. Our reference transcriptome was created with the Cellranger makeref command, and used genome assembly Mmul_10 as the reference genome and Ensembl v107 as the reference set of genomic annotations for the NHP samples, and Homo_sapiens.GRCh38 as the reference genome and Ensembl v107 as the reference for humans. In addition to the gene set from Ensembl, we added sequences for the two CAR transcripts used in the study for the NHP transcriptome.

For the scTCR-Seq libraries, we used the reference VDJ set created in [1] for NHPs, and the "refdata-cellranger-vdj-GRCm38-alts-ensembl-7.0.0.tar.gz" VDJ reference provided by 10x for humans.

**Initial QC and Filtering – NHP Data**

Fastqc was used to assess the quality of the sequencing, and the output metrics from Cellranger were used to assess the quality of the libraries. The "filtered_feature_bc_matrix.h5" file and "filtered_contig_annotations.csv" output files from Cellranger were combined into an AnnData object and loaded into scanpy[2]. scTCR-Seq for the validation NHPs was removed due to low quality. Since the libraries for R.315 and the libraries for R.301-R.304 were generated under different conditions, we decided to keep the two datasets separate rather than attempt to integrate them. Following the recommendations in Leucken and Theis[3], we analyzed various metrics of droplet quality to choose cutoffs for identifying cells. For the Initial NHP dataset, we retained cells that met the following criteria: <10% of the transcripts mapped to the mtDNA, genes detected > 900, and total transcripts between 2,500 and 25,000. For the validation dataset, we retained cells that met the following criteria: <10% of the transcripts mapped to the mtDNA, genes detected > 200, and total transcripts between 1000 and 25,000.

We then identified the 7,500 most variable genes in each of the two datasets and filtered the datasets to these genes along with *CD3E, TRAC, CD4, CD8A*, and the two CAR transcripts. We then applied PCA, and the neighbors, Leiden (clustering), and UMAP commands from Scanpy[2] with default parameters on each dataset and retained clusters enriched for *CD3E* and *TRAC*, and removed those enriched for monocyte and other non-T cell markers.

**Identification of Latent Space, Clusters, and Normalized Counts with scVI – NHP Data**

We then used scvi-tools (v0.19.0) to fit a variational autoencoder (VAE) to each of the datasets. For each dataset, we trained the variational autoencoder on our raw transcript counts using the parameters n_layers=2, n_latent=30, gene_likelihood="nb". We found a small monocyte cluster in the initial NHP dataset, removed these cells, and fit a new VAE to the data.

We used the “get_latent_representation” function from scVI to get the latent space from the VAE for each dataset. We then applied the neighbors, UMAP, and Leiden functions from Scanpy with default settings to obtain clusters and the UMAP dimensionality reduction for the two datasets. We used the “get_normalized_expression” function from scVI with library size set to 10,000 to obtain normalized counts for the cells.

**Identification of Cluster Markers with scVI**

To identify marker genes for each cluster, we applied the scVI function “differential_expression” with ‘group_by’ set to the initial clusters. We required cluster markers to meet the following criteria: 1) Bayes Factor > 3, 2) more than 10% of the cells in the cluster must express the gene, 3) lfc_median > 0.25, and 4) scVI’s FDR <0.05 must equal “TRUE”.

**Comparison of Immunological Signatures in NHP Data between CARPos and CARNeg cells**

We applied Decoupler’s WSUM (weighted sum) function to score each cell in the data by multiplying the normalized counts for each gene in each cell by +1 (upregulated) or -1 (downregulated) as indicated in the signature. We then performed a t-test to determine if the cell scores were different between the two groups.

**Identification of Bystander Signature in the Initial NHP**

We created a signature for bystander T cells by using scVI's differential expression function to identify marker genes for each cluster, and then applied additional filtering to limit the list to genes that were strongly upregulated / downregulated in Cluster 14 vs all other cells. Genes were considered upregulated in Cluster 14 if they: 1) were marked `is_de_fdr_0.05` by scVI, 2) had a Bayes Factor >3, 3) were expressed in at least 25% of the cells in the bystander cluster, and 4) had a median log2 fold change >1. Similarly, genes were considered downregulated in Cluster 14 if they 1) were marked `is_de_fdr_0.05` by scVI, 2) had a Bayes Factor >3, 3) were expressed in at least 25% of the non-bystander T cells, and 4) had a median log2 fold change < -1. The CAR transcript was excluded from this list.

In addition to this first gene signature: Signature 2: where we restricted ourselves to only CD8+ T Cells, Signature 3: where we only use $CAR^{Neg}$ CD8+ T Cells from the expansion timepoint. Finally, we manually created a fourth signature with a set of flow-detectable surface markers we identified for bystander T cells (weights set equal to +1).

**Assessment of Diversity and Overlap of T Cell Clones using scTCR-Seq**

We used the “vegan” package in R[4] which implements several metrics used for ecological studies. We measured diversity using Shannon diversity, and overlap using the Morisita Index.

**Initial QC and Filtering – Patient Data**

We followed the same steps described in the Initial QC and Filtering – NHP Data section. For the patient dataset, we retained cells that met the following criteria: <10% of the transcripts mapped to the mtDNA, genes detected >1,100, and total transcripts between 3,000 and 40,000.

**Identification of Latent Space, Clusters, and Normalized Counts with scVI – Patient Data**

We followed the same steps described in “Identification of Latent Space, Clusters, and Normalized Counts with scVI – NHP Data”.

**Identification of Bystander Cells in Patient Data**

For each rhesus gene in our bystander signatures, we identified the human ortholog. For cases where multiple rhesus genes mapped to the same human ortholog, we used the mean value of the rhesus log2FC's as its score. We then applied these signatures to the human data using Decoupler's WSUM function as described previously, and used a t-test with Welch correction to determine if a particular cluster had higher bystander scores than the remaining cells.


1 Gerdemann, U. *et al.* Identification and Tracking of Alloreactive T Cell Clones in Rhesus Macaques Through the RM-scTCR-Seq Platform. *Front Immunol* **12**, 804932 (2021). https://doi.org:10.3389/fimmu.2021.804932

2 Wolf, F. A., Angerer, P. & Theis, F. J. SCANPY: large-scale single-cell gene expression data analysis. *Genome Biol* **19**, 15 (2018). https://doi.org:10.1186/s13059-017-1382-0

3 Luecken, M. D. & Theis, F. J. Current best practices in single-cell RNA-seq analysis: a tutorial. *Mol Syst Biol* **15**, e8746 (2019). https://doi.org:10.15252/msb.20188746

4 vegan: Community Ecology Package. v. R package version 2.6-5 (2023).

Supplemental Figure 1

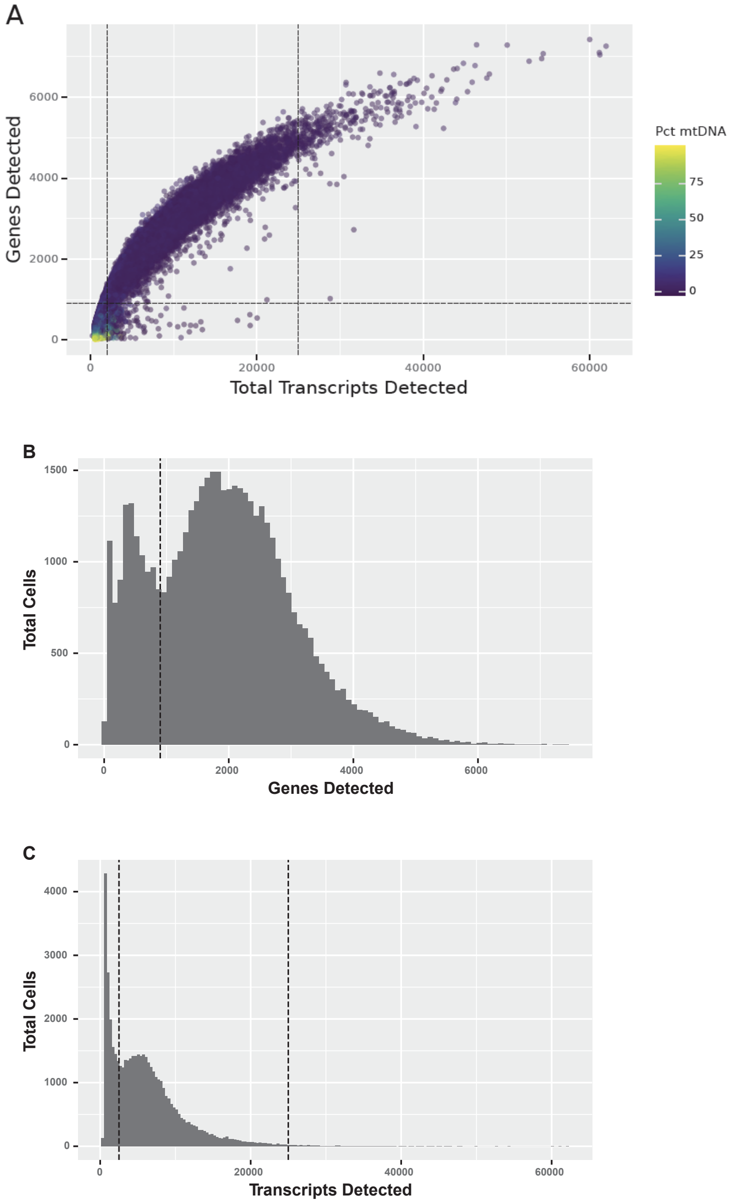


**Supplemental Figure 1 – Analysis of Droplet Quality Measures to Identify Cells for NHP analysis.**

(A) – Scatterplot of genes detected vs transcripts detected for each droplet, with share of reads mapping to mtDNA indicated by color of point. (B) Histogram of genes detected. (C) Histogram of transcripts detected. We retained cells with <10% of the transcripts mapped to the mtDNA, genes detected > 900, and total transcripts between 2,000 and 25,000, as indicated by the dashed lines.

**Supplemental Figure 2**

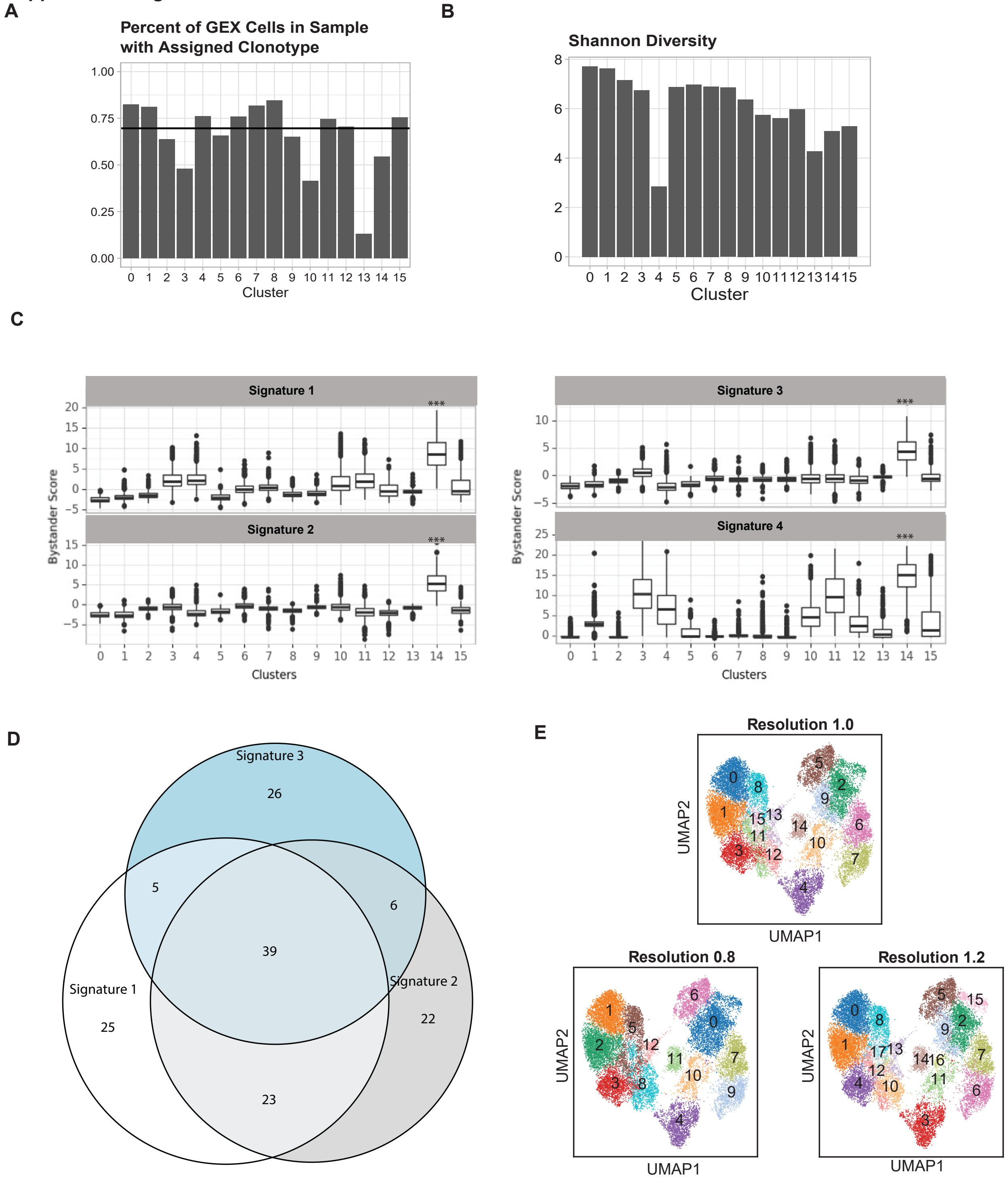

**Supplemental Figure 2 - Additional Quality Control Assessment of Initial NHP Dataset.**

(A) – Barplot displaying the percent of high quality GEX cells in each cluster that were assigned to a clonotype based on VDJ library. The percent of all high quality GEX cells assigned to a clonotype is 69.6%. (B) Shannon diversity of clones for each cluster. (C) Boxplot of signature scores for each cell in each cluster. (D) Venn diagram displaying number of shared genes among the three computationally determined signatures. (E) UMAP of clusters determined by three different resolutions.

**Supplemental Figure 3**

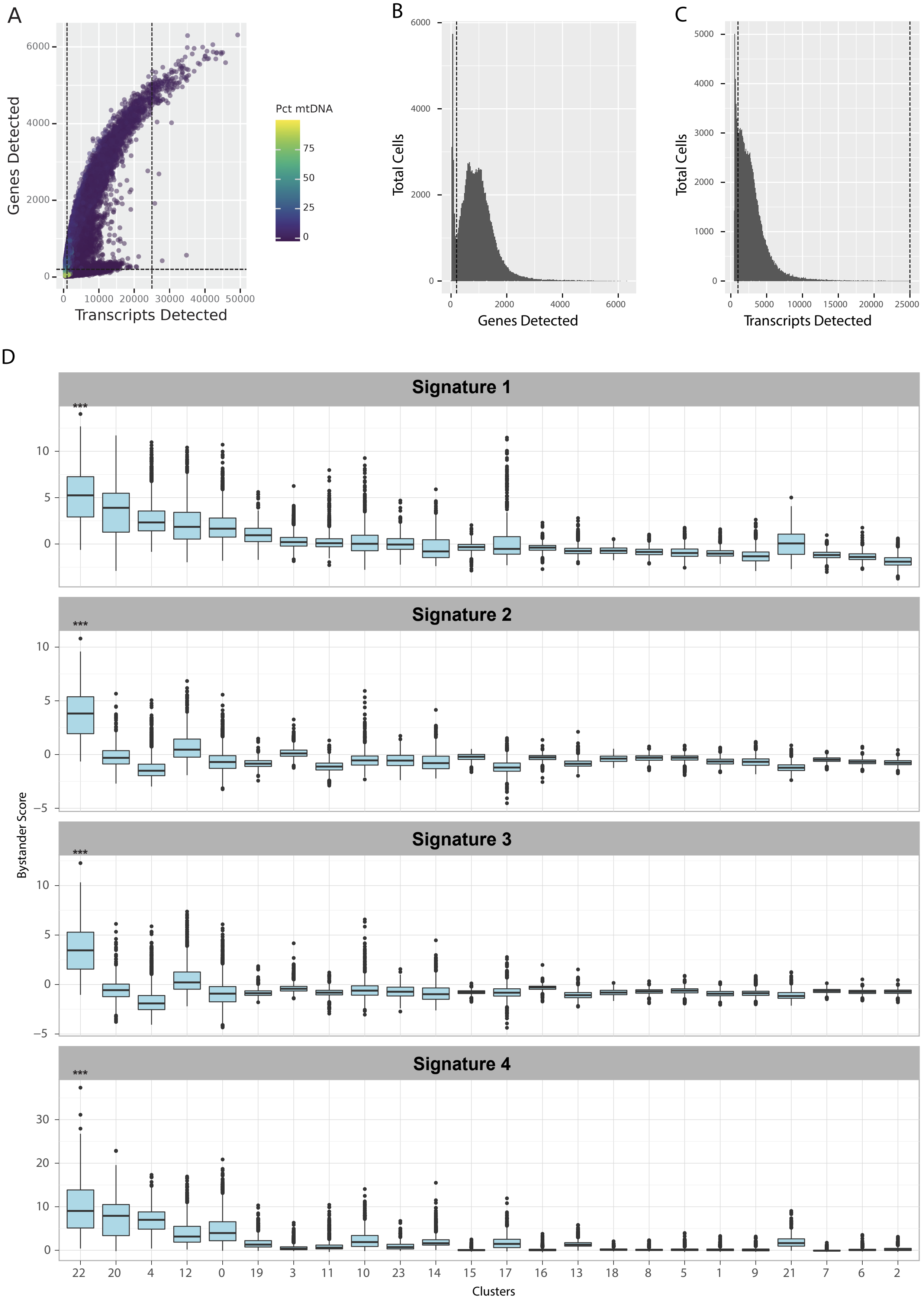

**Supplemental Figure 3 – Analysis of Droplet Quality Measures to Identify Cells, and Scores for All Bystander Signatures, on the Additional NHP Dataset.**

(A) – Scatterplot of genes detected vs transcripts detected for each droplet, with share of reads mapping to mtDNA indicated by color of point. (B) Histogram of genes detected. (C) Histogram of transcripts detected. We retained cells with <10% of the transcripts mapped to the mtDNA, genes detected > 200, and total transcripts between 1000 and 25,000. (D) Boxplot of signature cells for each cell in each cluster (*** = p-value< 0.001 for Welch-corrected t-test of signature score vs all other cells).

**Supplemental Figure 4**

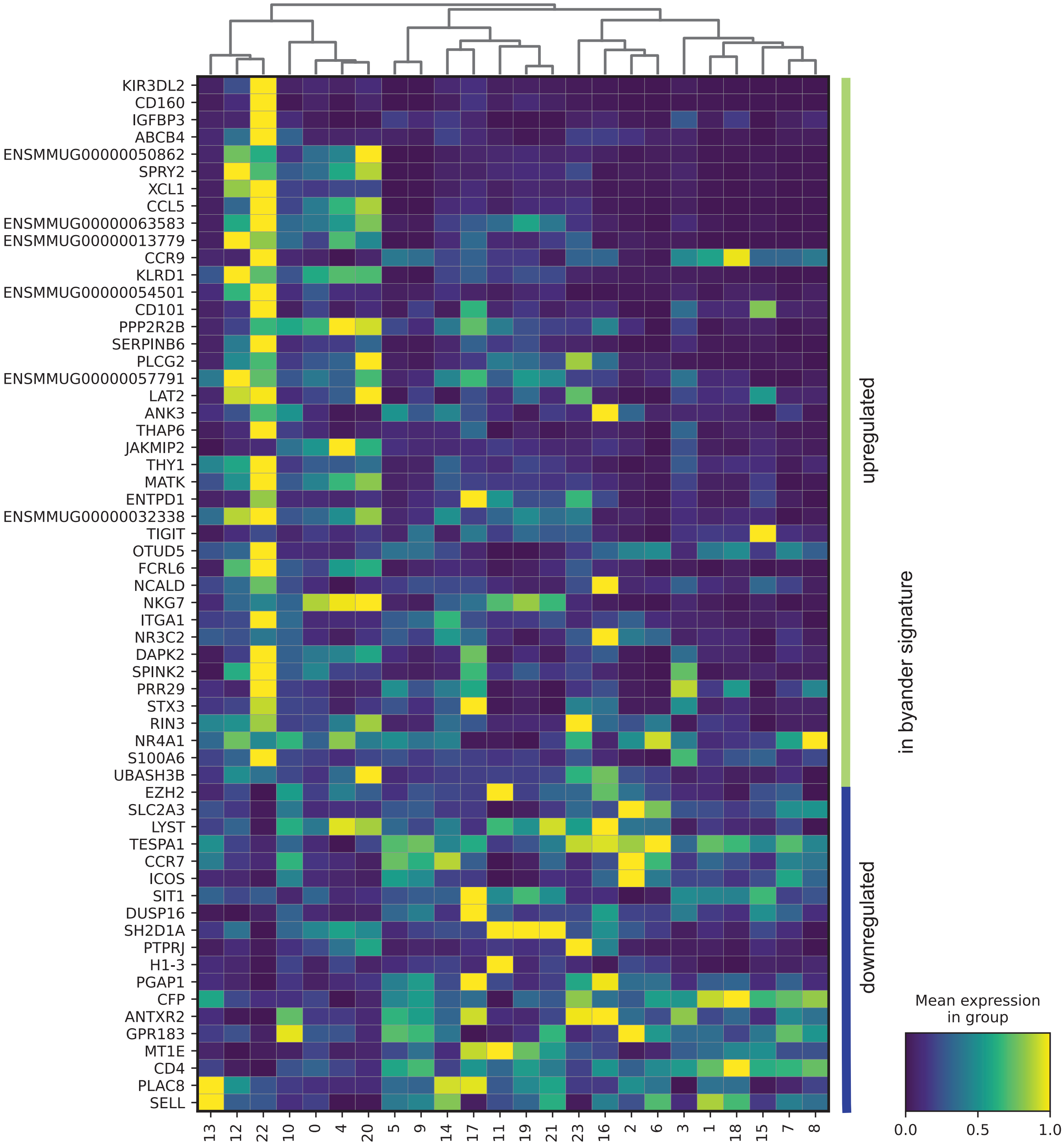

**Supplemental Figure 4 – Cluster 22 in the Additional NHP Dataset Expresses High Levels of Genes from Bystander Signature 1**

(A) Matrix plot of Signature 1 genes on each cluster in the Validation NHP Set. Each square shows the average normalized expressing of the given gene in a particular cluster. Values are row-scaled so that the cluster with highest expression has a score of 1 and the cluster with the lowest expression has a score of 0.

**Supplemental Figure 5**

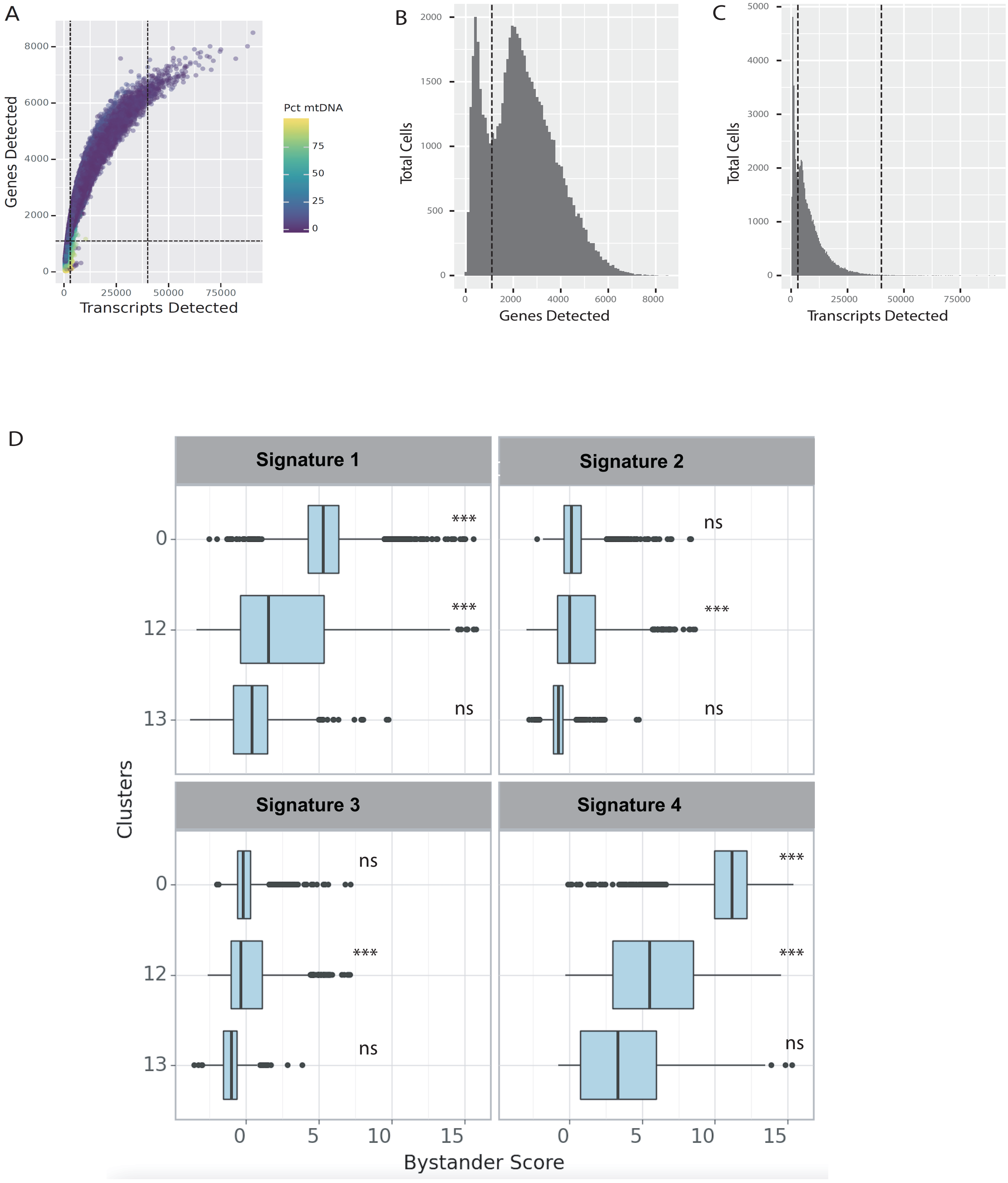

**Supplemental Figure 5 – Identification of Cutoffs for High Quality Cells, and Scores for All Bystander Signatures for the Patient Dataset.**

(A) – Scatterplot of genes detected vs transcripts detected for each droplet, with share of reads mapping to mtDNA indicated by color of point. (B) Histogram of genes detected. (C) Histogram of transcripts detected. We retained cells with <10% of the transcripts mapped to the mtDNA, genes detected > 1,100, and total transcripts between 3,000 and 40,000. (D) Boxplot of signature cells for each cell in each cluster on the dataset that was analyzed using a batch term for patient (*** = p-value< 0.001 for Welch-corrected t-test of signature score vs other CD8+ CAR- clusters, ns = not significant).

**Supplemental Figure 6**

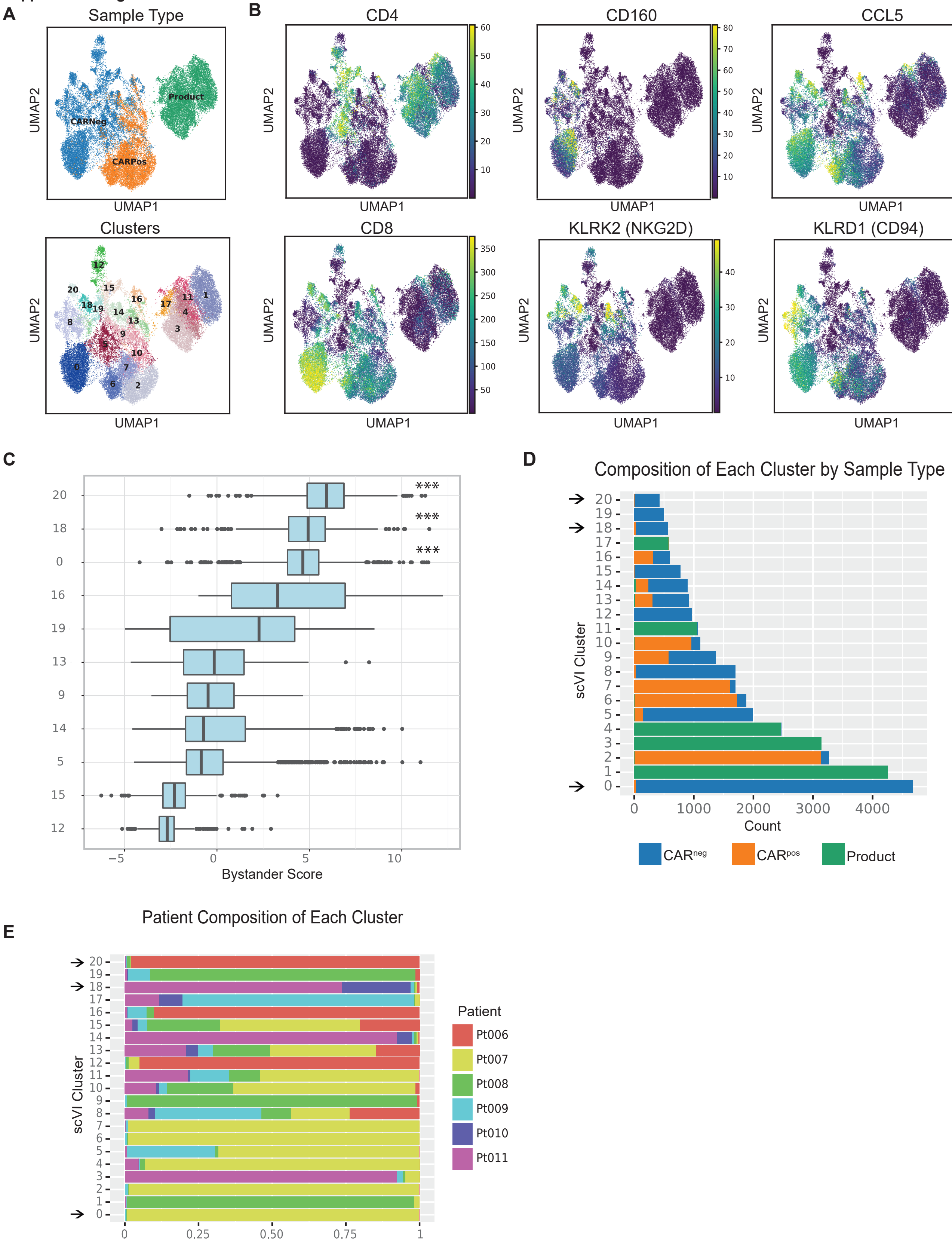

**Supplemental Figure 6 – Identification of Cutoffs for High Quality Cells, and Scores for All Bystander Signatures for the Patient Dataset (without Adjustment for Patient).**

(A) Plot of UMAP dimensionality reduction on patient dataset, colored by sample type and cluster respectively. (B) Plot of normalized expression for CD4 and key Bystander genes. (C) Boxplot of Bystander Signature 1 scores for each cell in each cluster (*** = p-value< 0.001 for Welch-corrected t-test of signature scores in cluster vs scores on other CD8+ CAR- T cells, ns = not significant). (D) Barplot displaying the composition of each cluster by sample type, with bystander clusters 0, 18, and 20 highlighted. (E) Barplot displaying the composition of each cluster by patient, with bystander clusters 0, 18, and 20 highlighted.

**Supplemental Tables**

**Table S1 - Maximum CAR T expansion NHP**

**Table S2 – Marker Genes for Initial NHP Clusters**

**Table S3 – Composition of Initial NHP Clusters by Timepoint, CAR Sort, and CAR Expression**

**Table S4 – Genes for Each Bystander Signature**

**Table S5 – Markers Genes for Clusters in the Additional NHP Dataset**

**Table S6 - Composition of Clusters in the Additional NHP Dataset, by Timepoint, Donor, CAR S Sort, and CAR Expression**

**Table S7 - Characteristics of Individuals in Patient Dataset**

**Table S8 - Marker Genes for Clusters in Patient Dataset (Patient-Adjusted Dataset)**

**Table S9 - Composition of Patient Dataset Clusters by Patient and Sample Type (Patient-Adjusted Dataset)**

**Table S10- Marker Genes for Clusters in Patient Dataset (Non-Adjusted Dataset)**

**Table S11 - Composition of Patient Dataset Clusters by Patient and Sample Type (Non-Adjusted Dataset)**

**Table S12 – Similarity of Bystander Cell Labeling in Both Patient Dataset Analyses**

**Table S13 – Percent of Patient Cells Labeled as Bystanders**

**Table S14 - Largest T cell Clones in Patient Dataset**

**Table S15 - TCR-CDR3 Identity in Patient Dataset**
